# Construction of a ceRNA network specific for granulosa cells in PCOS

**DOI:** 10.1101/2024.12.13.628409

**Authors:** E.G. Derevyanchuk, E.V. Butenko, T.P. Shkurat, L. Lipovich

## Abstract

**Background:** Polycystic ovary syndrome (PCOS) is a hormonal disease with a polygenic genetic model that occurs in 8 – 13% of women of reproductive age. In this study we constructed the granulose cell-specific competitive endogenous RNA (ceRNA) network in polycystic ovary syndrome. The ceRNA network analysis may provide novel insights into PCOS pathophysiology and identify potential therapeutic targets.

**Materials and methods:** The study included 6 women aged 26 to 34 years old: 3 women with PCOS and 3 women of the control group who underwent IVF procedure. Before RNA extraction GCs from three follicle samples of each group were collected and randomly pooled together as one mixed sample. Using RNA-sequencing we defined the microRNAs, lncRNAs and mRNAs expression profiles of GCs. The analysis of differential gene expression was performed using the DESeq v.1.39.0. The analysis of the interaction of microRNAs and lncRNAs was performed using RNAhybrid; of microRNAs and mRNAs - using the miRWalk database. Cytoscape 3.10.0 software was used to construct the ceRNA network.

**Results:** As a result of the differential gene expression analysis using the DESeq package we identified 3 differentially expressed microRNAs in PCOS, 132 - lncRNAs and 564 - mRNAs. The final ceRNA network included 3 microRNAs, 105 lncRNAs and 252 mRNAs.

**Conclusion:** This research underscores the importance of understanding ceRNA networks as a pathway for discovering biomarkers and developing treatments tailored to the unique challenges faced by individuals with PCOS. Further research is warranted to validate the identified interactions and explore their potential as diagnostic and therapeutic targets.

## Introduction

Polycystic ovary syndrome (PCOS) is a hormonal disease with a polygenic genetic model that occurs in 8 – 13% of women of reproductive age according to the statistics of the World Health Organization for 2023 (https://www.who.int/), and is one of the main causes of female infertility. It is often characterized by obesity and various metabolic disorders such as dyslipidemia, hyperinsulinemia and hyperandrogenism [1].

Despite numerous studies of pathogenesis, the mechanisms of PCOS occurrence remain unclear. A deeper understanding of the mechanisms underlying the development of PCOS is necessary to ensure early diagnosis and more effective treatment of the disease. Thanks to advances in the field of genetic and biological methods, it has been proven that long non-coding RNAs (lncRNAs) and microRNAs play a role in the occurrence and progression of PCOS and, thus, are diagnostic and therapeutic targets.

Molecules of lncRNAs have a length of more than 200 bases and do not encode proteins, since they do not have an open reading frame of sufficient length (although exceptions are described when a protein can be encoded [2]). It has been shown that lncRNAs regulate many processes, such as transcription, translation, cellular differentiation, regulation of gene expression and regulation of the cell cycle [3, 4]. It has been found that lncRNAs play a molecular role in the development of PCOS, mainly functioning as part of a competitive endogenous RNAs network, and correlate with some clinical phenotypes [5]. MicroRNAs are also important regulators of PCOS [6]. MicroRNAs are endogenously expressed RNA molecules with a length of 18 – 22 nucleotides that suppress gene expression at the post–transcriptional level by binding to the 3’-untranslated region of the target mRNA.

Currently, the most recognized is the competitive endogenous RNA network (ceRNA), in which lncRNA competes with protein-coding mRNAs for binding to microRNAs. It was found that ZFAS1 lncRNA can bind to miR-129, contributing to the expression of *HMGB1*, thereby affecting endocrine disorders, proliferation and apoptosis of ovarian granulosa cells in PCOS [7]. Granulosa cells (GC) are the most studied cell type in PCOS. As a rule, GCs are selected from women with or without PCOS undergoing in vitro fertilization to identify potentially pathogenic genes. A decrease in the level of MALAT1 in granulosa cells was revealed, which can contribute to the pathophysiological processes of PCOS by regulating the transmission of TGFß signals through the absorption of miR-125b and miR-203a [8]. PWNR2 plays an important role in the maturation of oocyte nuclei in PCOS, functioning as a ceRNA, reducing the availability of miR-92b-3p for binding to the TMEM120B target [9].

The aim of our study was to build a granulosa-specific competitive endogenous RNAs network and study its role in the emergence and development of PCOS. The analysis of the lncRNA/microRNA/mRNA network can serve as the basis for the treatment of patients with PCOS.

## Materials and methods

### Patient Recruitment

The study included 6 women aged 26 to 34 years old: 3 women with PCOS and 3 women of the control group who underwent IVF procedure at the Center for Human Reproduction and IVF (Rostov-on-Don, Russia) from 2021 to 2023. The main group consisted of women diagnosed with PCOS, which was confirmed in accordance with the Rotterdam criteria. The diagnosis was made if the patient had at least two of the following three criteria: 1) hyperandrogenism, confirmed clinically or biochemically, 2) oligo- or anovulation, 3) polycystic ovarian morphology, confirmed by ultrasound with the presence of > 12 follicles (2-9 mm in diameter) in each ovary or 20 antral follicles in both ovaries, or an enlarged ovary (> 10 ml). The control group consisted of women who underwent IVF procedures due to tubal factor infertility or male infertility. Endocrinological pathologies such as hyperprolactinemia, Cushing’s disease, adrenal hyperplasia or ovarian tumors were excluded from both groups. The objects for the study were samples of granulosa cells (GCs).

The research protocol was approved by the Bioethics Committee of the Academy of Biology and Biotechnology of the Southern Federal University. The study was performed in accordance with the Declaration of Helsinki [World Medical Association et al. World Medical Association Declaration of Helsinki: ethical principles for medical research involving human subjects //Jama. – 2013. – T. 310. – №. 20. – C. 2191-2194], and “Rules of clinical practice in the Russian Federation”, approved by the Ministry of Health of the Russian Federation (Order No. 266 dated June 19, 2003). All the participants signed the written informed consent.

### RNA-sequencing

Before RNA extraction GCs from three follicle samples of each group (3 PCOS and 3control) were collected and randomly pooled together as one mixed sample. Two libraries for sequencing were prepared from each mixed sample: 1 – for long RNA (coding and noncoding) sequencing; 2 – for microRNA sequencing. Libraries for long RNAs sequencing were prepared using NEBNext Ultra II RNA Library Prep kit for Illumina, rRNA depletion workflow was applied. Libraries for microRNAs sequencing were prepared using NEBNext Small RNA Library Prep Set for Illumina. RNA sequencing of long and small RNA libraries was conducted on the Miseq (Illumina) platform.

### Bioinformatics analysis

Mapping and counting the number of reads using STAR 2.7.9a with the outFilterMismatchNmax 3 parameter was performed. Gene expression was quantified using RPKM (Reads Per Kilo bases per Million reads).

The analysis of differential gene expression (DEGs) was performed using the DESeq v.1.39.0 package [10] in R. The change in gene expression in GCs of women with PCOS was evaluated. RNAs with a log2 fold change (log2 (FC)) >2 and p <0.05 were considered upregulated in PCOS compared to control samples. RNAs with log2 (FC) <−2 and p <0.05 were determined as downregulated. It is considered that the genes with more than 15 reads in the sequencing results are expressed, while the genes below 15 reads are not expressed.

The genes encoding mRNAs, lncRNAs, and microRNAs were classified using the corresponding Ensembl identifiers.

Gene set enrichment analysis (GSEA) using GO and KEGG ontologies was carried out using an R package clusterProfiler with FDR<0.05 for each ontology.

The analysis of the interaction of microRNAs and lncRNAs after removal of microRNAs not characterized in miRBase and homologous sequences was performed using RNAhybrid (Linux version) [11]. The analysis of the interaction of microRNAs and mRNAs was carried out using the miRWalk database [12] with a restriction on the score/bindingp parameter > 0.95.

Finally, Cytoscape 3.10.0 software [13] was used to construct the ceRNA network from the differentially expressed miRNAs, mRNAs, and lncRNAs in PCOS patients.

The programming language R v.4.2.3 was used for calculations.

## Results

In this study, using RNA-sequencing we defined the microRNAs, lncRNAs and mRNAs expression profiles of GC isolated from women with PCOS and healthy women. As a result of the differential gene expression analysis using the DESeq package we identified 3 differentially expressed microRNAs (expression of microRNAs hsa-miR-652-3p and hsa-miR-193b-3p was significantly increased (p < 0.05) in PCOS, whereas microRNA hsa-miR-181a-2-3p - lowered), 132 - lncRNAs and 564 - mRNAs.

We conducted the analysis of gene set enrichment (GSEA) using GO (Gene Ontology) and KEGG (Kyoto Encyclopedia of Genes and Genomes) ontologies. According to the results of the GO analysis (Figure 1), among the genes with differential expression included in the network, the largest number of genes encode cell adhesion proteins; control the response to signaling molecules; belong to the genes of the signaling pathway of cell surface receptors; genes associated with the processes of control and modulation of signaling pathways in the cell; genes associated with the processes of controlling and modulating the ways in which cells communicate with each other and with the environment; genes encoding the processes of cell movement.

**Figure 1.**
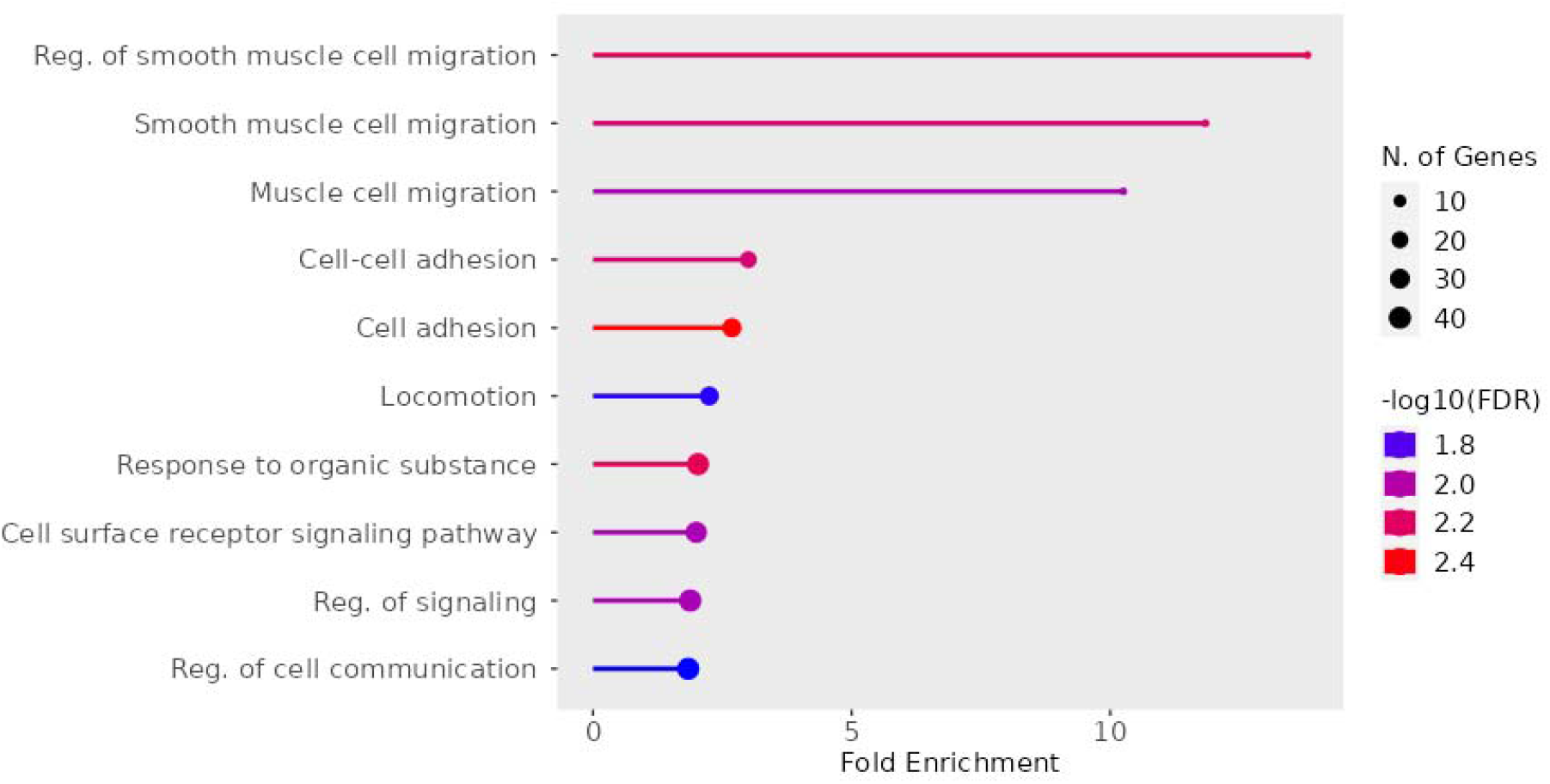
The main biological processes in GC controlled by genes in the ceRNA network associated with PCOS, according to the results of GO analysis.

The presence of the “Proteoglycans in cancer” signaling pathway in the KEGG analysis (Figure 2) indicates a potential link between the processes involved in the regulation of proteoglycans and the pathophysiology of PCOS. PCOS may show altered expression of proteoglycans in the ovaries or other tissues. This can lead to impaired cell adhesion, changes in the intercellular matrix and impaired ovarian function. At the same time, the analysis of our data using KEGG showed (Figure 2) the presence of the signaling pathway “Interaction between the extracellular matrix and receptors” (ECM-receptor interaction), which indicates a potential disruption in communication between ovarian cells and their extracellular environment. One possible explanation is that in the ovaries with PCOS, there may be a change in the composition and structure of the ECM. This may include changes in the amount and type of collagens, proteoglycans, and other ECM components. Such changes affect cell adhesion, proliferation, differentiation and migration of ovarian cells, which can lead to disorders of folliculogenesis and ovulation. The interaction between ECM receptors and ECM components triggers various intracellular signaling pathways that play an important role in regulating the growth, differentiation and function of ovarian cells. In PCOS, these signaling pathways can be disrupted, leading to ovarian dysfunction.

**Figure 2.**
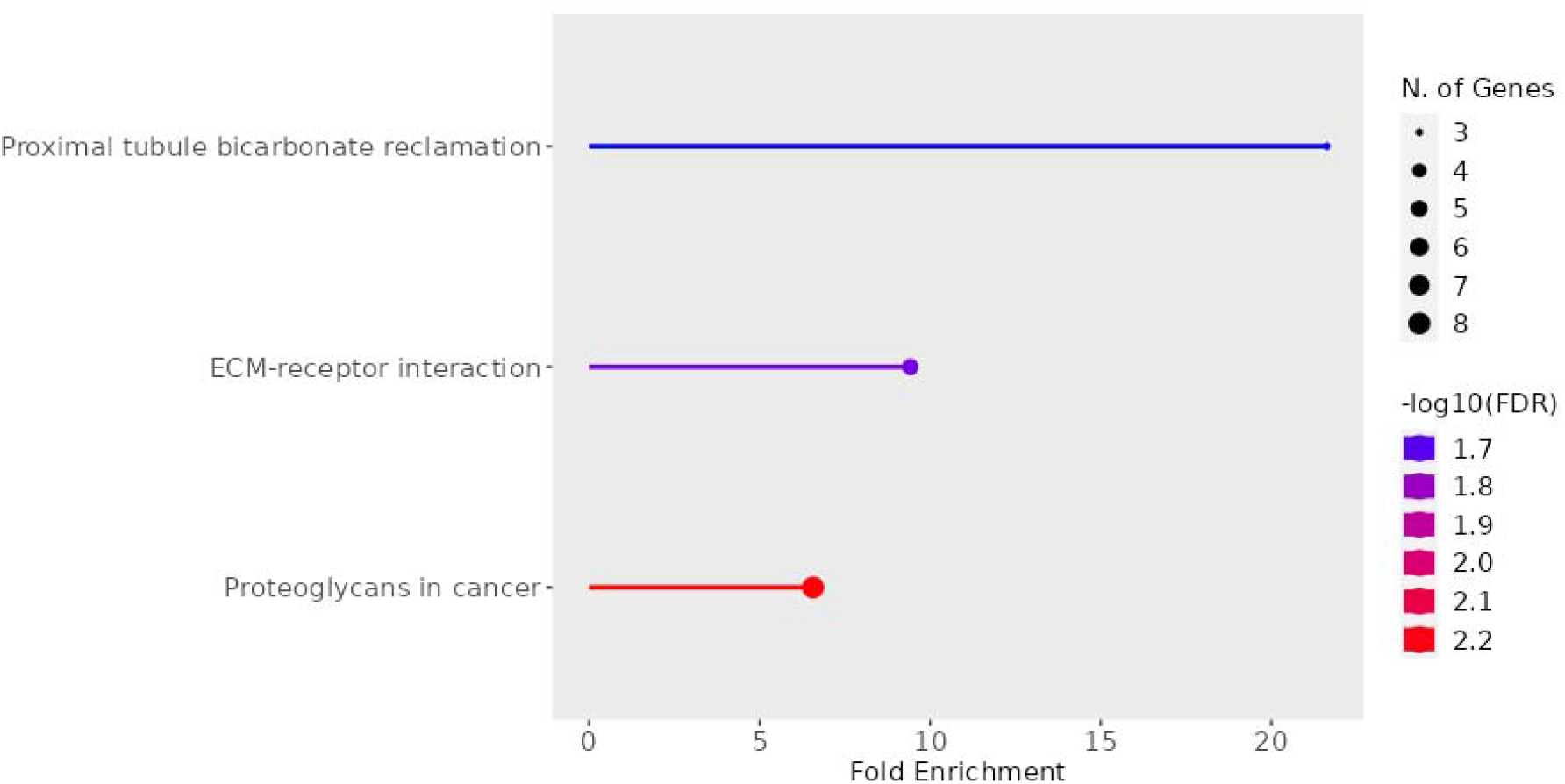
The main biological processes in GC controlled by genes in the ceRNA network associated with PCOS, according to the results of KEGG analysis.

After removal of miRBase-uncharacterized microRNAs and homologous sequences, 3 mature microRNA sequences (hsa-miR-652-3p, hsa-miR-181a-2-3p, hsa-miR-193b-3p) were included in the analysis. As a result of the analysis of the interaction of differentially expressed microRNAs and lncRNAs in granulosa cells using RNAhybrid, 105 interaction events were identified. RNAhybrid is a tool for determining the minimum free energy during hybridization of long and short RNA [12]. Hybridization is performed in a kind of domain mode, i.e. the short sequence hybridizes with the most appropriate part of the long one. This tool is primarily designed to predict targets for microRNAs.

The lncRNAs included in the interaction analysis (33) are able to bind to all three microRNAs: BAIAP2-DT; BET1-AS1; CAMTA1-AS2; ERVK9-11; GABPB1-AS1; GAS6-AS1; GDNF-AS1; GLIDR; GRIK1-AS1; H19; HS1BP3-IT1; IL6R-AS1; KCNMA1-AS3; LERFS; LINC00707; LINC01291; LINC01355; LINC01502; LINC01659; LINC01807; LINC02019; LINC02145; LINC02186; LINC02299; LINC02732; LINC02830; NCAM1-AS1; PAX8-AS1; PHEX-AS1; PLAC4; SLC9A3-AS1; SPATA13-AS1; ST8SIA6-AS1; ZNF571-AS1, except GABRG3-AS1 lncRNA which has binding sites only in hsa-miR-181a-2-3p and hsa-miR-193b-3p.

miRWalk is an open source platform providing an intuitive interface that generates predicted and validated microRNA binding sites of known human, mouse, rat, dog and cow genes [13]. miRWalk is based on the prediction of the microRNA target site using the TarPmiR software based on the random forest method to search for the complete sequence of transcripts, including 5’-UTR, CDS and 3’-UTR. In addition, it integrates results from other databases with predicted and confirmed target microRNA (mRNA) interactions. We identified 160 unique interaction events for 138 mRNAs in the miRWalk database for three microRNA sequences (hsa-miR-652-3p, hsa-miR-181a-2-3p, hsa-miR-193b-3p). Moreover, one mRNA target (*PDE4D*) had binding sites to all three microRNAs. The remaining mRNA targets possessed binding sites for only two of the three differentially expressed microRNAs in granulosa cells in PCOS.

We combined the microRNAs - lncRNAs and microRNA - mRNA pairs to reconstruct the lncRNA—miRNA—mRNA regulatory network in PCOS: 3 microRNAs, 105 lncRNAs and 252 mRNAs were used to construct the ceRNA network. The ceRNA network was visualized with the Cytoscape program using the yOrganic layout (Figure 3).

**Figure 3.**
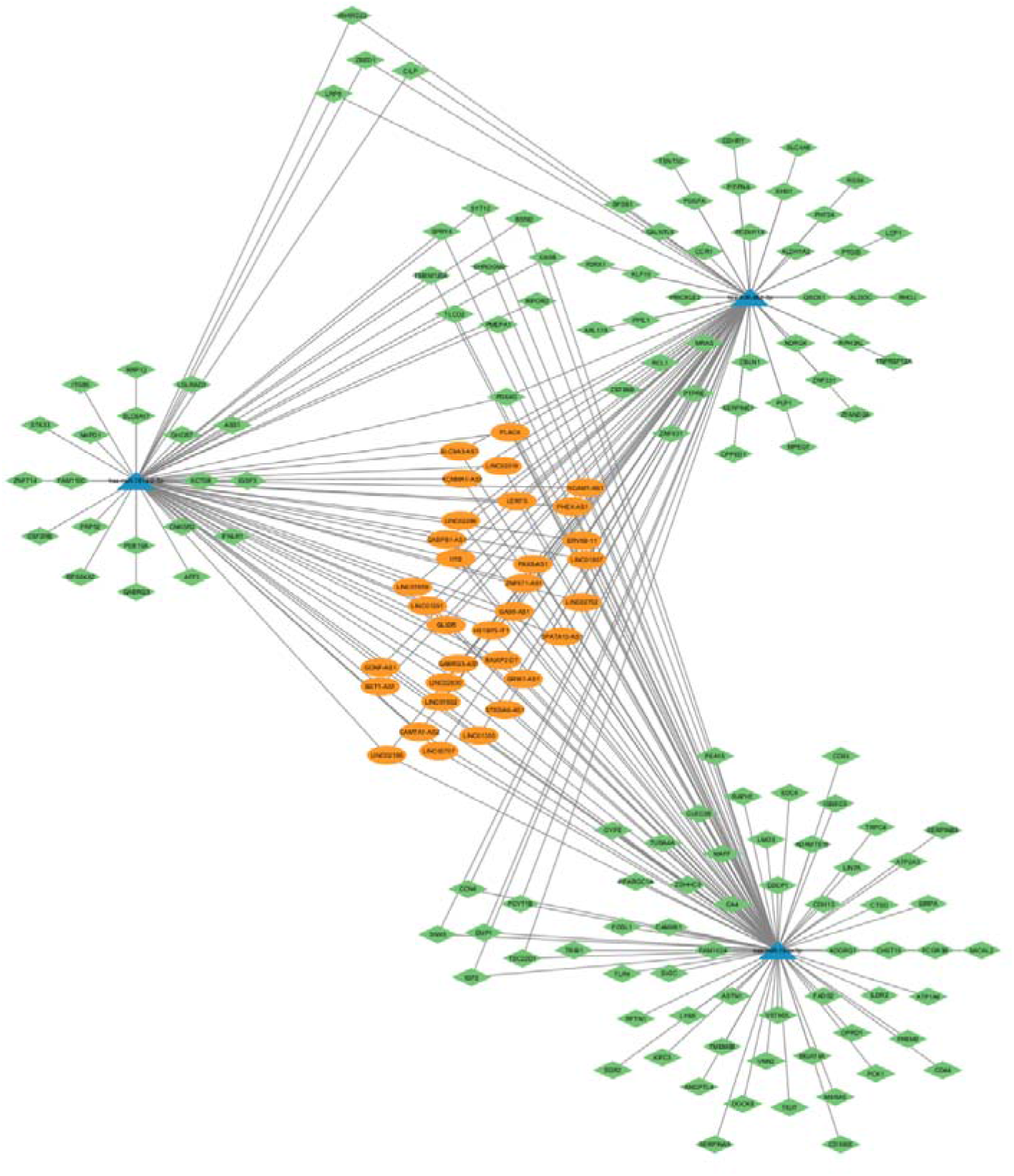
CeRNA network, specific for GC in PCOS. Blue triangles – microRNAs; green rhombuses – mRNAs; orange ovals – lncRNAs

LncRNAs are known to play a role in gene expression regulation because they can partially inhibit microRNA activity. They bind to microRNAs like a sponge and act as a competing endogenous RNA, regulating the function of the target microRNA, thereby indirectly affecting mRNA levels.

## Discussion

PCOS is a common endocrine disease of the reproductive system and a serious problem for women of childbearing age. PCOS affects both female fertility and many organs in the body [14]. Currently, there is no effective drug for the complete cure of PCOS, and many patients suffer from it all their lives. The pathogenesis of polycystic ovary syndrome is complex, and an in-depth study of the molecular mechanism underlying it will help to identify targets for its diagnosis and treatment.

In our previous study we developed a pipeline for ceRNA network inference in PCOS, using public PCOS and control transcriptome data [15]. In that study we identified 75 genes protein–coding genes (23 with increased expression, 52 – with reduced), 8 lncRNA genes (1 with increased expression, 7 – with reduced) and 1 miRNA with reduced expression (hsa-mir-3681) which were significantly differentially expressed between PCOS and control groups. Gene set enrichment analysis (GSEA) with the GO and KEGG ontologies showed that the differentially expressed genes form a pro-inflammatory signature associated with cytokine activity and the performance of the corresponding receptors. The miRNA and lncRNA interactions were not detected, but significant expression of hsa-miR-3681-5p in granulosa cells in polycystic ovaries was revealed and specific ceRNA network of hsa-miR-3681-5p with protein-coding genes involved in cell proliferation and apoptosis (CCDC69; DCLK1); intercellular adhesion (CEACAM6); transmembrane transport (SV2C); anti-inflammatory response (CXCL6; LAIR1) and modulation of cholesterol metabolism (HMGCR) was constructed.

In this study we conducted RNA-sequencing analysis of GCs from women with PCOS and healthy controls which revealed significant alterations in the expression profiles of microRNAs, lncRNAs, and mRNAs. This finding aligns with previous studies demonstrating the crucial role of non-coding RNAs in PCOS pathogenesis [16]. The identification of three differentially expressed microRNAs – hsa-miR-652-3p and hsa-miR-193b-3p showing increased expression, and hsa-miR-181a-2-3p exhibiting decreased expression – in PCOS GCs warrants further investigation. The altered expression of these microRNAs suggests their involvement in the complex molecular mechanisms underlying PCOS. For example, miR-652-3p has been implicated in various cellular processes, including cell proliferation and apoptosis [17], while miR-181a-2-3p is known to regulate immune responses [18]. The functional consequences of the observed changes in their expression levels within the context of PCOS remain to be elucidated. The substantial number of differentially expressed lncRNAs and mRNAs highlights the widespread transcriptional dysregulation characteristic of PCOS GCs. The GO and KEGG enrichment analyses, although not detailed in the provided results, are crucial for determining the biological pathways and cellular components affected by these differentially expressed genes. These analyses should illuminate the functional consequences of the transcriptional changes, potentially revealing key processes perturbed in PCOS, such as steroidogenesis, cell cycle regulation, and inflammatory responses.

While this observation underscores the importance of these interactions in PCOS pathogenesis, further investigation is needed to determine the specific functional consequences of these interactions and their impact on downstream target genes. The limited sample size necessitates validation in larger cohorts to strengthen these findings and explore the clinical relevance of these interactions.

## Conclusion

This study establishes a foundation for understanding the role of ceRNA network in PCOS pathogenesis. As a result of the analysis of interactions between target mRNAs and microRNAs, microRNAs and lncRNAs differentially expressed in granulosa cells in PCOS, the final network of competitive endogenous RNAs included 3 microRNAs, 105 lncRNAs and 252 mRNAs. Further research is warranted to validate the identified interactions and explore their potential as diagnostic and therapeutic targets. Targeting ceRNA network may offer promising avenues for improving PCOS diagnosis, prevention, and treatment.

## Author contributions

**PhD, Assoc. Prof. Derevyanchuk E.G. (corresponding author):** Performed data analysis, bioinformatic analysis, figure preparation, wrote the initial draft.

**PhD, Assoc. Prof. Butenko E.V**.: Was primarily responsible for data collection and sample preparation. Also contributed significantly to data interpretation and manuscript revisions.

**Dr**., **Prof. Shkurat T.P**.: Led the study design, provided critical review of the manuscript.

**Dr**., **Prof. Lipovich L**.: Provided valuable input on the experimental design and interpretation of results. Contributed to manuscript revisions.

